# A TPR scaffold couples signal detection to OdhI phosphorylation in metabolic control by the protein kinase PknG

**DOI:** 10.1101/2021.06.11.448168

**Authors:** María-Natalia Lisa, Adrià Sogues, Nathalie Barilone, Meike Baumgart, Magdalena Gil, Martín Graña, Rosario Durán, Ricardo M. Biondi, Marco Bellinzoni, Michael Bott, Pedro M. Alzari

## Abstract

Signal transduction is essential for bacteria to adapt to changing environmental conditions. Among many forms of post-translational modifications, reversible protein phosphorylation has evolved as a ubiquitous molecular mechanism of protein regulation in response to specific stimuli. The Ser/Thr protein kinase PknG modulates the fate of intracellular glutamate by controlling the phosphorylation status of the 2-oxoglutarate dehydrogenase regulator OdhI, a function that is conserved among diverse actinobacteria. PknG has a modular organization characterized by the presence of regulatory domains surrounding the catalytic domain. Here we present an investigation through *in vivo* experiments as well as biochemical and structural methods of the molecular bases of the regulation of PknG from *C. glutamicum* (*Cg*PknG), in the light of previous knowledge available for the kinase from *M. tuberculosis* (*Mtb*PknG). We found that OdhI phosphorylation by *Cg*PknG is regulated by a conserved mechanism that depends on a C-terminal domain composed of tetratricopeptide repeats (TPR) essential for metabolic homeostasis. Furthermore, we identified a conserved structural motif that physically connects the TPR domain and a flexible N-terminal extension of the kinase that is involved in docking interactions with OdhI. Based on our results and previous reports, we propose a model in which the TPR domain of PknG couples signal detection to the specific phosphorylation of OdhI. Overall, the available data indicate that conserved PknG domains in distant actinobacteria retain their roles in kinase regulation in response to nutrient availability.

**IMPORTANCE:** Bacteria control the metabolic processes by which they obtain nutrients and energy in order to adapt to the environment. In this way, the metabolic characteristics of a microorganism determine its ecological role and its usefulness in industrial processes. Here, we use genetic, biochemical, and structural approaches to study a key component in a system that regulates glutamate production in *C. glutamicum*, a species that is used for the industrial production of amino acids. We elucidated molecular mechanisms involved in metabolic control in *C. glutamicum*, which are conserved in related pathogenic bacteria. The findings have broader significance for diverse actinobacteria, including microorganisms that cause disease as well as environmental species used to produce billions of dollars of amino acids and antibiotics every year.

## INTRODUCTION

The large and ancient bacterial phylum *Actinobacteria* comprises species with very diverse lifestyles and physiological adaptations, including soil inhabitants, pathogens as well as plant or animal commensals (1). The eukaryotic-like Ser/Thr protein kinase (STPK) PknG and its FHA (ForkHead-Associated) substrate OdhI (Oxoglutarate dehydrogenase Inhibitor) are at the core of a conserved signal transduction pathway that modulates central metabolism in distant actinobacteria. Both in *Corynebacterium glutamicum*, a soil bacterium used for the industrial production of amino acids, as well as in the pathogen *Mycobacterium tuberculosis*, PknG modulates the 2-oxoglutarate dehydrogenase activity in the Krebs cycle (2–4) by controlling the phosphorylation status of the regulator OdhI (called GarA in the genus *Mycobacterium*) (2–5). Biochemical studies have demonstrated that unphosphorylated OdhI/GarA inhibits the E1 component (OdhA) of the 2-oxoglutarate dehydrogenase complex whereas this inhibition is relieved by OdhI/GarA phosphorylation by PknG (2–4, 6, 7). Moreover, early studies for the two species revealed that *pknG* disruption leads to an accumulation of intracellular glutamate (2, 8), pointing out that PknG acts by promoting catabolism at the expense of 2-oxoglutarate usage in nitrogen assimilation. On top of this, it was recently found that PknG senses the availability of amino-donor amino acids to control metabolism and virulence in *M. tuberculosis* (9–11). These findings have received much attention (10), as a deeper understanding of PknG regulation can be instrumental for downstream applications in the biotech and pharmaceutical areas.

PknG has a unique modular organization characterized by the ubiquitous presence of a flexible N-terminal segment and a C-terminal domain composed of tetratricopeptide repeats (TPR) flanking the kinase catalytic core (12–14). An additional rubredoxin (Rdx)-like domain occurs immediately adjacent to the catalytic core in PknG from mycobacteria and most other actinobacteria but not in corynebacteria (2). Previous structural studies of PknG have focused on the protein from *M. tuberculosis* (*Mtb*PknG) (12, 13). We have shown that the N-terminal extension and the TPR domain of *Mtb*PknG regulate the selectivity for GarA without significantly affecting the intrinsic kinase activity, whereas the Rdx domain downregulates catalysis by limiting access to a profound substrate-binding site (13). Rdx domains are known to transmit redox stimuli and, consistent with this, evidence has been reported pointing out that perturbations of the metal center in PknG lead to alterations of the kinase activity (15). However, relatively little is known about the regulatory mechanisms of PknG isoforms that lack an Rdx domain.

The gene *pknG* is found within a conserved operon that contains two other genes, *glnX* and *glnH*, which encode a putative transmembrane protein and a putative glutamine-binding lipoprotein, respectively (2, 11). The observation that disruption of any of those genes in *C. glutamicum* led to a similar phenotype consisting of a growth defect in medium containing glutamine as the sole carbon source (2) suggested a common role of the protein products in metabolic homeostasis. Supporting this early hypothesis, evidence has been recently reported that, in mycobacteria, PknG and GlnX are functionally linked and that GlnH specifically binds amino acids able to stimulate GarA phosphorylation by the kinase (11). This led to the proposal that GlnH senses amino acid availability within the bacterial periplasm and transmits this information across the membrane *via* GlnX to activate PknG by protein-protein interactions (11). Most interesting, a PknG truncation mutant lacking the TPR domain failed to restore the growth defect of a *pknG*-disrupted mycobacterial strain, suggesting that this domain, often involved in protein-protein interactions (16), mediates molecular associations required for the kinase function (11).

To investigate the conservation of mechanisms involved in the regulation of PknG, we studied the kinase isoform from *C. glutamicum* (*Cg*PknG), which is devoid of an Rdx domain. We provide evidence that the C-terminal region of *Cg*PknG, bearing the TPR domain, is crucial for the efficient phosphorylation of OdhI and for the kinase function in metabolic homeostasis. Moreover, our results point out that the recruitment of the FHA substrate is regulated by a conserved phosphorylation-dependent mechanism regardless of the absence of an Rdx domain. Finally, by comparing three high-resolution crystal structures of *Cg*PknG and an available structure of *Mtb*PknG (12), we identified a conserved motif able to link the N-terminal extension and the TPR domain. Interestingly, the evidence suggests that the Rdx domain, absent in corynebacteria, and the TPR domain would constitute independent regulatory mechanisms. Overall, our results indicate that common PknG domains in distant actinobacteria share similar functions in kinase regulation, linking PknG to the control of central metabolism in response to nutrient availability.

## RESULTS

### The C-terminal region of *Cg*PknG is required for phosphorylation events that modulate metabolism

To investigate the domains required for the function of *Cg*PknG, we employed a previously characterized *C. glutamicum* Δ*pknG* mutant strain able to grow in rich medium but unable to grow in medium containing glutamine as the sole carbon source (2). *Cg*PknG domain boundaries were defined based on a previous characterization of *Mtb*PknG (13) (47% amino acid identity), and plasmids were designed for the expression of *Cg*PknG truncation mutants (Fig. 1A) in *C. glutamicum* Δ*pknG* using the endogenous gene promoter. All strains grew normally in medium containing glucose and all versions of the kinase were detected by Western-blot (Fig. S1).

**Figure 1.**
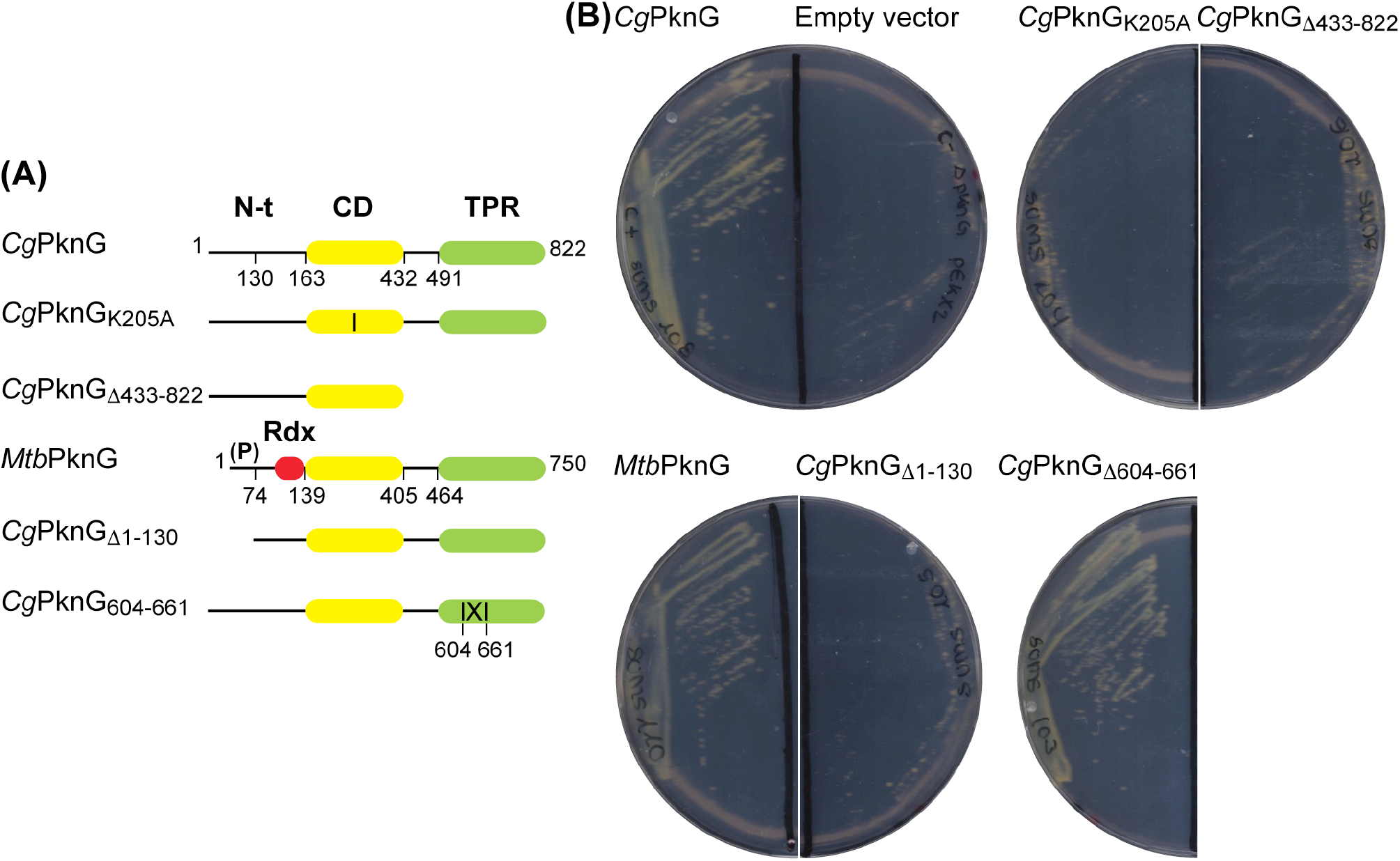
Complementation of *C. glutamicum* Δ*pknG* (2) with different PknG versions. (A) Schematic representation of the kinase variants tested in complementation assays in this study. The structured domains of the protein are shown as colored rectangles: the Rdx domain in red, the catalytic domain (CD) in yellow, and the TPR domain in green. The vertical line in the CD of mutant *Cg*PknG_K205A_ represents the amino acid substitution. The (P) symbol indicates the cluster of autophosphorylation sites in the N-terminal region (N-t) of *Mtb*PknG (13). The IXI symbol in the TPR domain of *Cg*PknG_604-661_ represents an internal segment of deleted amino acids. (B) Complementation of the *ΔpknG* strain with different *pknG* versions. Complementation was assessed by growth on CGXII plates with 100 mM glutamine as sole carbon source after 3 days at 30°C. PknG variants capable to complement the *ΔpknG* strain were *Cg*PknG, *Mtb*PknG and *Cg*PknG_Δ604-661_. The empty pEKEx2 vector was used as a negative control.

In contrast to wild type *Cg*PknG, the mutant *Cg*PknG_K205A_, which harbors a substitution of the invariant catalytic lysine, did not complement the growth defect of *C. glutamicum* Δ*pknG* on glutamine (Fig. 1B), indicating that the kinase activity is required for protein function. Additionally, a *Cg*PknG truncation mutant lacking residues 433-822 was unable to restore bacterial growth on glutamine, pointing out, in agreement with previous results for *Mtb*PknG (11), that the region of *Cg*PknG located C-terminally to the catalytic core is necessary for the kinase role in the control of metabolism. Moreover, *Mtb*PknG did complement the growth defect of *C. glutamicum* Δ*pknG*, stressing the functional conservation between distant kinase isoforms. A *Cg*PknG deletion mutant devoid of residues 1-130 failed to restore the growth of *C. glutamicum* Δ*pknG* on glutamine, however the low amount detected of this kinase version precludes drawing conclusions from this observation. Together, these results support a conserved requirement of the C-terminal region of PknG for phosphorylation events that modulate metabolism in response to amino acid availability.

### A conserved phosphorylation-dependent mechanism for substrate recruitment

To investigate the molecular mechanisms of metabolic control by the kinase activity of *Cg*PknG, we first tested the ability of recombinant *Cg*PknG to phosphorylate OdhI and GarA *in vitro. Cg*PknG phosphorylated OdhI and GarA to a similar extent (Fig. 2A), confirming the ability of *Cg*PknG to phosphorylate the FHA substrate and evidencing that structural differences between OdhI and GarA (4, 17), either in the FHA domain or in the N-terminal phosphorylatable region, do not influence the kinase activity. Moreover, *Cg*PknG phosphorylated GarA in the same peptide as *Mtb*PknG (3) (Fig. S2), equivalent to the OdhI peptide phosphorylated by *Cg*PknG (2).

**Figure 2.**
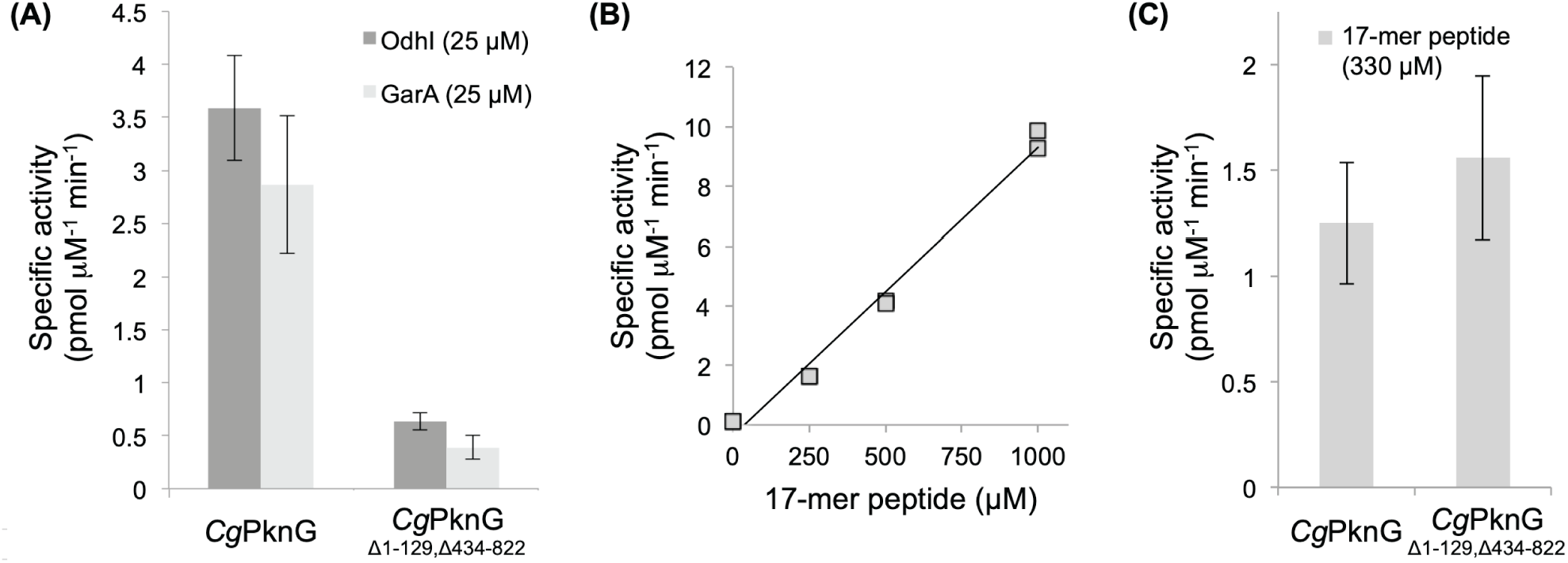
*Cg*PknG and CgPknG_Δ1-129, Δ434-822_ and their relative kinase activities. (A) Relative kinase activities of *Cg*PknG and CgPknG_Δ1-129, Δ434-822_ against OdhI and GarA. (B) Kinase activity of *Cg*PknG for different concentrations of the 17-mer peptide substrate SDEVTV**ETTS**VFRADFL. (C) Relative kinase activity of *Cg*PknG and CgPknG_Δ1-129, Δ434-822_ against the 17-mer peptide. Measurements were performed at least twice; error bars represent the scattering among average values obtained in independent determinations.

The N-terminal segment of *Mtb*PknG contains auto-phosphorylation sites (Thr23, Thr32, Thr63 and Thr64) (3) (Fig. 1A and Fig. S3) that act as essential anchoring points for the recruitment of GarA by interacting with the pThr-binding FHA domain of the regulator (3, 13). Despite the crucial role of the N-terminal extension of the kinase in substrate selectivity, its primary structure is poorly conserved. Therefore, to determine whether or not the role of the kinase N-terminal segment in the recruitment of the FHA substrate is conserved in spite of sequence divergence, we first investigated the auto-phosphorylation of *Cg*PknG. Despite no phosphorylation was detected in the purified recombinant protein, four phosphorylation sites (Thr14, Thr68, Thr92 and Thr93) were identified by mass spectrometry within the N-terminal extension of *Cg*PknG after incubating the kinase with ATP and Mn(II) (Fig. S3 and Fig. S4).

Next, we studied the ability of *Cg*PknG to phosphorylate a substrate lacking an FHA domain, using for this the previously reported 17-mer SDEVTV**ETTS**VFRADFL peptide (13) centered around the phosphorylatable ETTS motif that is conserved among OdhI/GarA homologs (2). The kinase activity of *Cg*PknG varied linearly with the concentration of the 17-mer peptide up to 1 mM, indicating a high *K*_M_ (> 1 mM) and the slope providing a measure of the catalytic efficiency (*k*_cat_/*K*_M_) of (9.0 ± 0.4) 10^−3^ pmol µM^-2^ min^-1^ for this substrate (Fig. 2B). By comparison, the phosphorylation of OdhI by *Cg*PknG was approximately 3-fold higher than for the 17-mer peptide even though a *ca*. 15-fold lower concentration of OdhI was used (Fig. 2A and Fig. 2C), indicating a *ca*. 45-fold higher activity towards OdhI due to the FHA domain acting as a kinase docking site.

Finally, we tested the kinase activity of a *Cg*PknG deletion mutant lacking residues 1-129 and 434-822. *Cg*PknG_Δ1-129,Δ434-822_ displayed a *ca*. 7-fold lower activity against OdhI compared to the full-length enzyme, whereas phosphorylation of the 17-mer substrate was unaffected (Fig. 2A and Fig. 2C). These results indicate that neither residues 1-129 within the N-terminal extension, nor the TPR domain of *Cg*PknG had an effect on the intrinsic kinase activity, supporting previous evidence for *Mtb*PknG (13) that both regions contribute to stabilize the enzyme-FHA substrate complex.

Overall, our results indicate that diverse PknG isoforms recruit the FHA substrate OdhI (or GarA) *via* a conserved phosphorylation-dependent mechanism.

### A conserved overall topology

To investigate the structural basis of the regulation of a PknG isoform lacking an Rdx domain, we solved a high-resolution crystal structure of *Cg*PknG_ΔN-t_ (see below) in complex with the non-hydrolysable ATP analog AMP-PNP (Table 1). The final atomic model contains two copies of *Cg*PknG within the asymmetric unit, encompassing residues 123-799 and 125-798, respectively, including a short fragment of the N-terminal segment (hence the name *Cg*PknG_ΔN-t_), the kinase catalytic core and the TPR domain (Fig. 3 and Fig. S5A). The protein is monomeric, consistent with analytical ultracentrifugation that did not provide evidence in favor of *Cg*PknG dimerization (Fig. S5B), similar to previous results for *Mtb*PknG (13). Additionally, *mFo–DFc* sigma-A-weighted electron density maps clearly revealed the bound nucleotide and two Mg(II) atoms at the active site of each *Cg*PknG molecule. Notably, even though we used full-length *Cg*PknG in our crystallization assays, we found no evidence for residues 1-122 in electron density maps. Edman degradation experiments revealed that the N-terminal residue of crystallized *Cg*PknG was Val123, suggesting that the kinase N-terminal segment was partially degraded during crystal growth and that, as similarly reported for *Mtb*PknG (12), it is probably unstructured in most of its length.

**Table 1.**
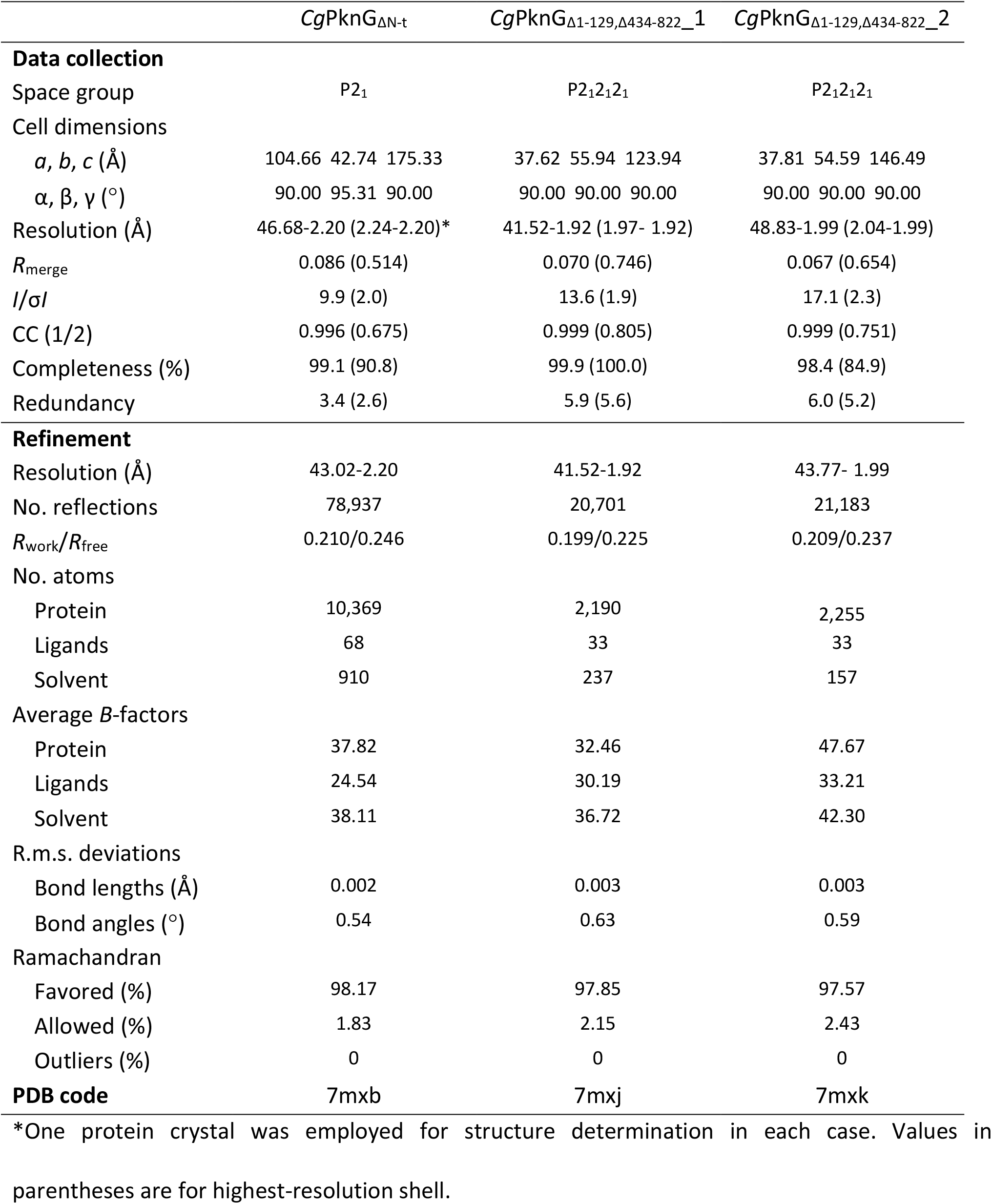
Data collection and refinement statistics.

**Figure 3.**
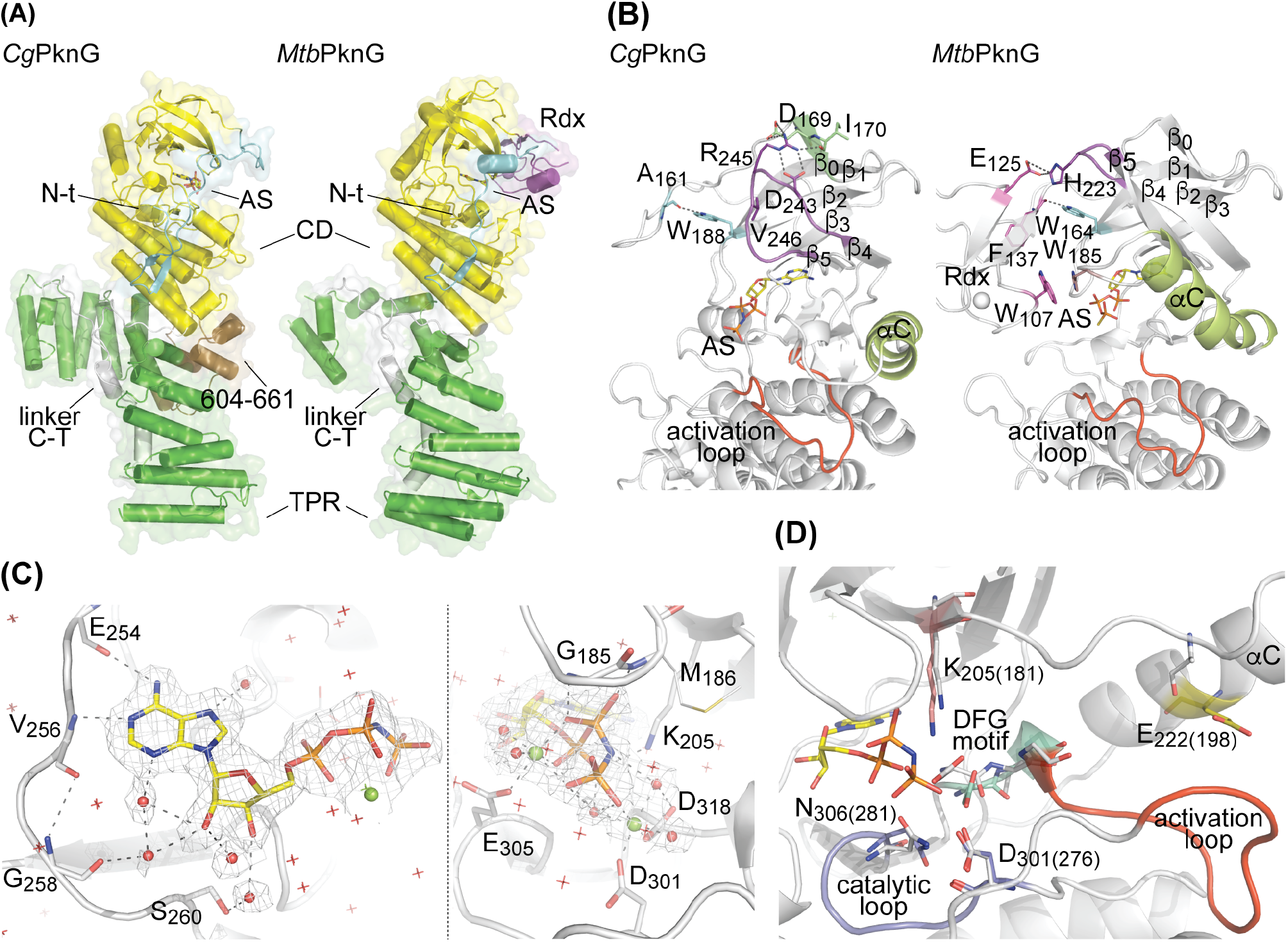
The crystal structure of *Cg*PknG_ΔN-t_. (A) Comparison of *Cg*PknG_ΔN-t_ and *Mtb*PknG_Δ1-73_ (12) (PDB code 2PZI). The chain A in each crystal structure is shown (RMSD of 2.35 Å for 532 aligned residues). The non-hydrolysable ATP analog AMP-PNP bound to the active site (AS) of *Cg*PknG_ΔN-t_ is depicted in sticks. N-t: N-terminal region; CD: catalytic domain; linker C-T: linker between the catalytic domain and the TPR domain. (B) Comparison of *Cg*PknG_ΔN-t_ and *Mtb*PknG_Δ1-73, Δ406-750_ (13) (PDB code 4Y12). The highlighted kinase domain residues or motifs adopt distinct conformations in the absence or in the presence of an Rdx domain. (C) The ATP binding site of *Cg*PknG _ΔN-t_ with a bound AMP-PNP molecule. The AMP-PNP molecule and the protein residues interacting with it are shown in sticks. Water molecules are depicted as red spheres or stars and Mg(II) atoms are shown as green spheres. The *2mFo–DFc* electron density is contoured to 1.0 σ and presented as a mesh. Dashed lines represent atomic interactions. (D) Functionally important and conserved residues within the kinase active site are shown for *Cg*PknG_ΔN-t_. Gray sticks correspond to residues in *Mtb*PknG_Δ1-73, Δ406-750_ (13) (PDB code 4Y12), numbered between brackets.

*Cg*PknG and *Mtb*PknG (12) share the same overall fold and topology, except for the absence of a regulatory Rdx domain in *Cg*PknG that leads to a more accessible active site (Fig. 3A). As expected, kinase domain residues or motifs involved in contacts with the Rdx domain in *Mtb*PknG (12, 13) adopt distinct conformations in *Cg*PknG (Fig. 3B). Residue Trp188 in *Cg*PknG (equivalent to Trp164 in *Mtb*PknG), located in the β_2_ strand and adjacent to the G-rich loop, interacts with the N-terminal segment. The loop connecting strands β4 and β5 (loop β4-β5) is found in *Cg*PknG in close association with the kinase N-lobe, with residue Val246 (His223 in *Mtb*PknG) buried within a pocket and residues Asp243 and Arg245 in contact with the strand β0. Besides, the helix αC does not interact with strands β4 and β5 and its C-terminal tip is displaced, in *Cg*PknG compared to *Mtb*PknG, towards the kinase activation loop.

Regardless of these differences, nucleotide binding within the active site of *Cg*PknG parallels the previous description for *Mtb*PknG (13) (Fig. 3C), consistent with a conserved set of residues within the ATP binding site region of the kinase. Also similar to *Mtb*PknG (12, 13), most functionally important and conserved motifs in the active site of *Cg*PknG exhibit conformations compatible with a standard eukaryotic protein kinase active state, and the activation loop is stabilized in an open and extended conformation, permissive for substrate binding in the absence of phosphorylation (Fig. 3D). Nevertheless, *Cg*PknG residue Glu222 is found away from the catalytic Lys205, pointing out of the active site due to an outward conformation of the helix αC, as previously reported for *Mtb*PknG (12, 13).

Compared to *Mtb*PknG, *Cg*PknG contains an additional motif (residues 604-661) in the TPR domain, adjacent to the catalytic core (Fig. 1 and Fig. 3A). However, a *Cg*PknG truncation mutant lacking residues 604-661 did complement the growth defect of *C. glutamicum* Δ*pknG* on glutamine, suggesting that this motif is not crucial for the kinase function.

### A conserved motif connects the N-terminal segment and the TPR domain

The TPR domain of *Mtb*PknG influences the FHA substrate selectivity and we have previously proposed that this depends on the stabilization of a β-hairpin in the N-terminal extension of the kinase (13). In spite of sequence divergence, this secondary structure motif is conserved in *Cg*PknG (Fig. 4A). Both in *Cg*PknG and *Mtb*PknG the N-terminal β-hairpin is stabilized by interactions with the catalytic core and the linker between this and the TPR domain (linker C-T, see also Figs. 1A and 3A). Notably, the linker C-T simultaneously contacts the N-terminal segment, the catalytic core and the TPR domain of the kinase. To explore the significance of such interactions, we solved the high-resolution crystal structures of the truncation mutant *Cg*PknG_Δ1-129,Δ434-822_ in two different isoforms (Table 1). According to the electron density maps, the N-terminal β-hairpin was not stabilized in any of the structures of *Cg*PknG_Δ1-129,Δ434-822_ (Fig. 4B), suggesting that this motif is responsive to the C-terminal region of the kinase. These results indicate that the linker C-T physically connects the conserved N- and C-terminal regions flanking the kinase catalytic core.

**Figure 4.**
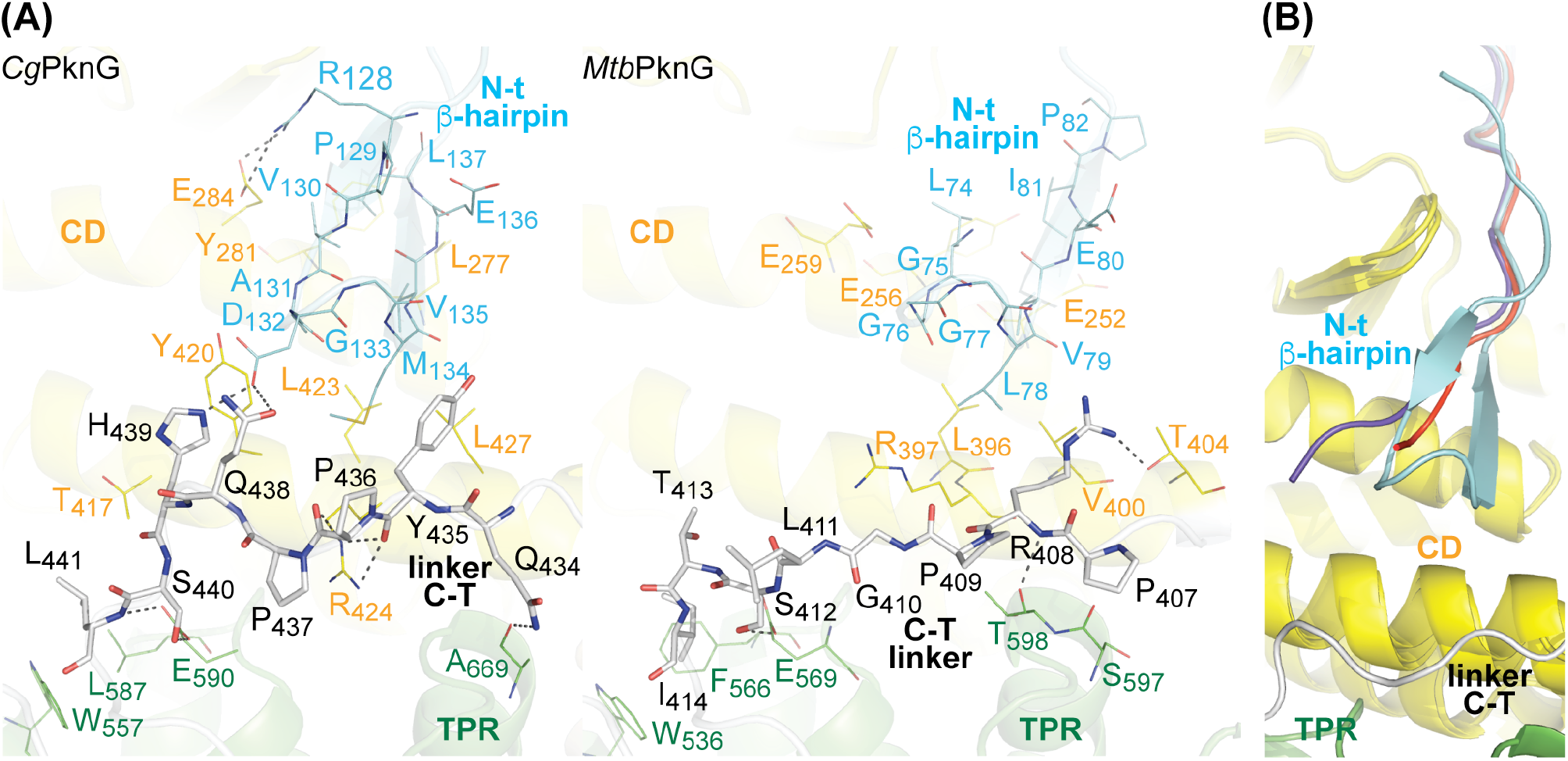
The linker C-T simultaneously interacts with an N-terminal β-hairpin, the catalytic core and the TPR domain of PknG. (A) Comparison of the crystal structures of *Cg*PknG_ΔN-t_ (this work) and *Mtb*PknG_Δ1-73_ (12) (PDB code 2PZI). The chain A in each crystal structure is shown. Selected residues within the linker C-T are shown in sticks. Residues conforming the N-terminal β-hairpin are depicted as lines. Residues of the catalytic core or the TPR domain involved in polar or hydrophobic interactions with the N-terminal β-hairpin or the linker C-T are also shown as lines. Dashed lines represent polar interactions. (B) The crystal structures of *Cg*PknG_ΔN-t_ and *Cg*PknG_Δ1-129,Δ434-822_ are superimposed. The RMSD values between the chain A in the structure of *Cg*PknG_ΔN-t_ and the structures of *Cg*PknG_Δ1-129,Δ434-822_ are 1.04 Å and 0.79 Å for 282 and 288 aligned residues, respectively. The N-terminal extension of *Cg*PknG_Δ1-129,Δ434-822_ is colored in blue or red.

## DISCUSSION

The phosphorylation-dependent stabilization of enzyme-substrate complexes is a widespread mechanism among STPKs that enables the efficient phosphorylation of specific cellular targets (18). PknG controls metabolism in corynebacteria and mycobacteria by modulating the phosphorylation status of the FHA regulator OdhI (or GarA) (2, 9), a task that requires the N-terminal extension of the kinase. Despite the relatively high sequence divergence of this segment, it has a roughly conserved distribution of charged amino acids, Pro and Gly residues in diverse species (Fig. S3), and comprises auto-phosphorylation sites both in *Cg*PknG and in *Mtb*PknG (3) (Fig. S3 and Fig. S4). The N-terminal extension of PknG is dispensable for the phosphorylation of a surrogate peptide lacking an FHA domain (Fig. 2C and (13)) and, conversely, the presence of the FHA domain in OdhI or GarA enables a much more efficient phosphorylation by full-length PknG (Fig. 2A, Fig. 2C and (13)). Overall, our results support a conserved phosphorylation-dependent mechanism for the recruitment of the FHA substrate *via* the kinase N-terminal extension.

Kinase domain motifs that play regulatory roles in eukaryotic protein kinases (ePKs) adopt different conformations in PknG isoforms depending on the presence or the absence of an Rdx domain. In *Cg*PknG the loop β4-β5 fills the pocket formed by the β-sheet in the kinase N-lobe, whereas this loop is exposed to the solvent in *Mtb*PknG (12, 13) (Fig. 3B). The pocket and the motifs that may fill it (*i*.*e*., the N-lobe cap) lay on top of the catalytic Lys and are features associated with the regulation of ePKs (19). Besides, the helix αC, an important regulatory motif in ePKs (20, 21), is displaced in *Cg*PknG towards the kinase activation loop when compared to *Mtb*PknG (12, 13) (Fig. 3B). Consistent with previous findings for ePKs (22), the crystal structures of both *Cg*PknG and *Mtb*PknG (12, 13) exhibit relatively high B-factors for the loop β3-αC and the N-terminal end of the helix αC, indicating that this motif is highly dynamic. Interestingly, while the Rdx domain in *Mtb*PknG restraints the position of the helix αC by interacting with the loop β3-αC (12, 13), the position adopted by the helix αC in *Cg*PknG generates a pocket that is reminiscent of the PIF-pocket in AGC kinases (22, 23) (Fig. S6). However, irrespective of the structural differences noted between *Cg*PknG and *Mtb*PknG (13), in both kinase isoforms the ATP phosphates are properly positioned in the active site despite the absence of a salt bridge between the conserved Glu in the helix αC and the catalytic Lys, while other conserved catalytically relevant motifs exhibit conformations compatible with an ePK active state (20) (Fig. 3C and Fig. 3D). Thus far, there is no evidence revealing regulatory mechanisms that depend exclusively on motifs within the kinase catalytic domain. The Rdx module of *Mtb*PknG (absent in *Cg*PknG) remains the sole regulatory element known to modulate the intrinsic activity of PknG (13, 15). It is worth noting that Rdx-mediated regulation appears to act independently of the modulation of substrate specificity by FHA-mediated docking interactions.

As the assembly of new domain combinations into complex proteins is linked to speciation and segregation into distinct phylogenetic groups (24, 25), we performed a phylogenetic analysis of PknG orthologs to seek for hints about the PknG-Rdx association (Fig. S7). In line with such notion, PknG orthologs, distinguished by their unique domain organization, are broadly distributed within *Actinobacteria* and also mostly restricted to this bacterial phylum. A homologue of *Mtb*PknG is, however, found in *Ktedonobacter racemifer*. This Gram-positive spore-forming bacterium belongs to *Chloroflexi* and grows in filamentous colonies similarly to a number of actinobacteria (26). *Chloroflexi* is an ancient phylum proposed to be at or very close to the root of the bacterial phylogenetic tree (27). Besides, a readily detectable homologue of PknG from *K. racemifer* is that from *Calothrix* sp. from the ancient phylum *Cyanobacteria*. The fact that both of these PknG homologues harbor an Rdx domain (defined by the presence of a PknG_rubred Pfam PF16919 domain or two CxxCG motifs) suggests that such domain architecture either preceded the evolution of *Actinobacteria*, being then differentially lost in some lineages, or that the gene of an Rdx-containing PknG homolog was horizontally transferred to *Chloroflexi* and *Cyanobacteria*. We favor the former, more parsimonious hypothesis because several non-actinobacterial ancient sequences include an Rdx domain whereas the genus *Corynebacterium* lacks the module. It remains enigmatic why the Rdx domain was lost in evolution in this genus.

The overall topology of PknG is conserved irrespective of the presence or the absence of an Rdx domain (Fig. 3A). The relative position of the TPR and the catalytic domain of *Cg*PknG is similar to that of *Mtb*PknG (12). Compared to the mycobacterial isoform, *Cg*PknG contains an additional, intriguing motif (residues 604-661) in the TPR domain (Fig. 1A and Fig. 3A) that increases its interface with the catalytic core. However, according to our *in vivo* tests, such motif is not essential for the role of *Cg*PknG in metabolic homeostasis (Fig. 1B). In contrast, the C-terminal region of *Cg*PknG (residues 433-822) was required for complementing the *C. glutamicum* Δ*pknG* mutant strain (Fig. 1B), replicating previous results for *Mtb*PknG (11) and pointing to a conserved role of the TPR domain in signal transduction. Notably, in *Cg*PknG as in *Mtb*PknG the linker C-T bridges the N-terminal segment and the TPR domain (Fig. 4A and (12)), both regions involved in the regulation of the kinase selectivity for the FHA substrate (13). The linker C-T is stabilized by conserved interactions with residues along the concave surface of the TPR domain (Fig. 3A and (12)). According to a recent proposal (11), this surface might constitute a binding site for GlnX, so that the transduction of extracellular stimuli would imply a conformational change of the linker C-T from its position in the free form of the kinase.

Taking together the available evidence, we propose that the TPR domain of PknG functions as a localization scaffold that, by mediating an interaction between the kinase and the transmembrane protein GlnX, transduces a signal about amino acid availability detected by GlnH (Fig. 5). The PknG-GlnX interaction likely produces a conformational change in the linker C-T, which couples the detection of the signal to the specific recruitment of the FHA substrate *via* the N-terminal segment of the kinase. Given that the specific set of multidomain proteins in genomes sets constraint on the topology of pathways and networks that carry out regulatory processes (28), the co-occurrence of *pknG, glnX, glnH* and *odhI* in actinobacteria (2, 11), together with the functional links found among the respective proteins, therefore suggests the conservation of the associated molecular mechanism that evolved in this phylum to control metabolism in response to nutrient availability.

**Figure 5.**
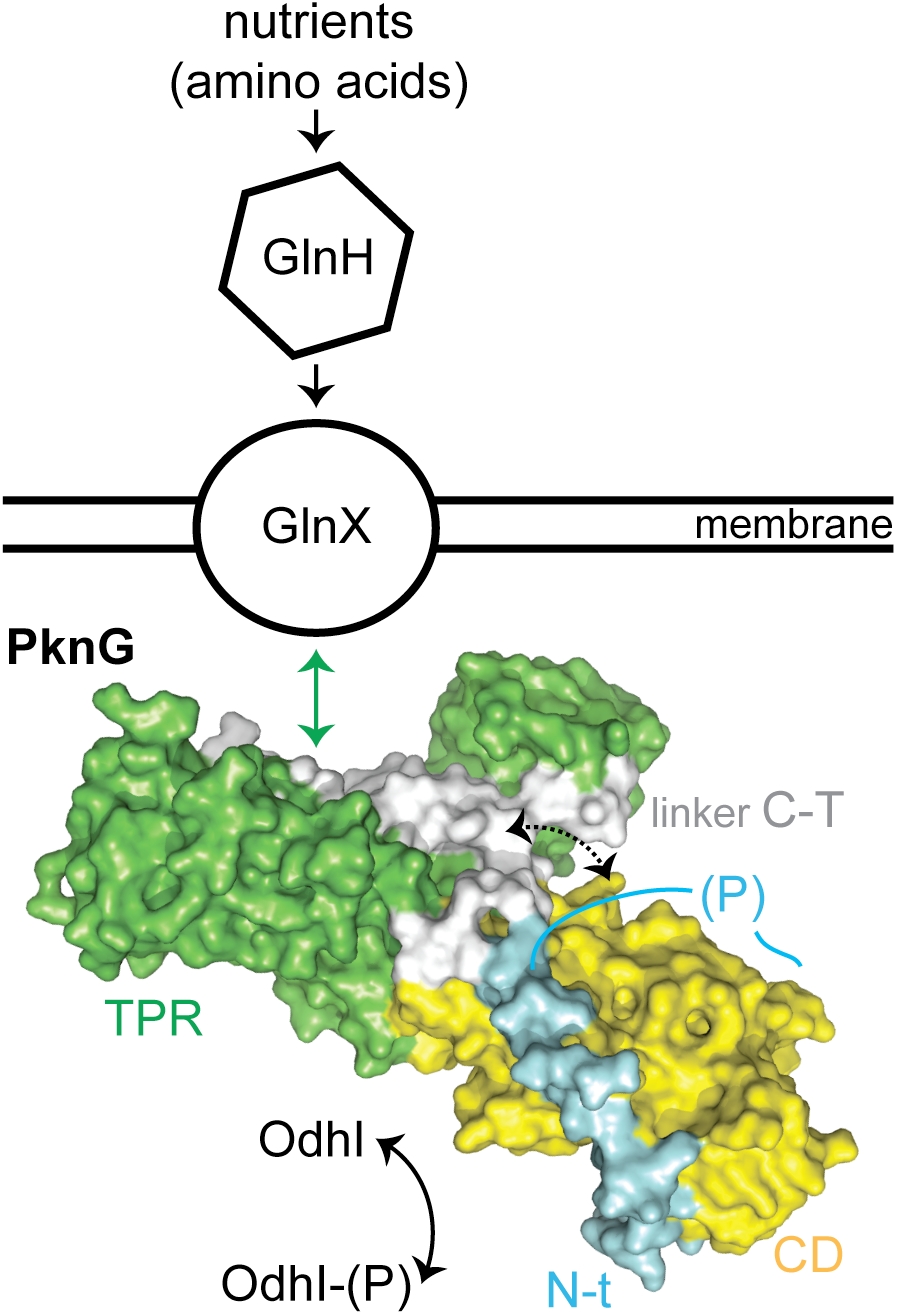
Proposed model for the role of the TPR domain in the *Cg*PknG function. The available genetic, biochemical, and structural evidence suggests that the TPR domain might act as a localization scaffold that, providing a surface for the interaction between the kinase and the transmembrane protein GlnX, would couple signal detection to OdhI phosphorylation by modulating the conformation of the linker C-T.

## MATERIALS AND METHODS

### Complementation assays

All plasmids used in this study are listed in Table 2. Plasmids for complementation assays were generated by Genscript (Leiden, The Netherlands) from the previously described pEKEx2-*pknG*_St_ template plasmid (2). The *C. glutamicum* Δ*pknG* strain (2) was transformed with each of the plasmids carrying the relevant *pknG* variants, or with the pEKEx2 vector lacking an insert, as previously described (29). Then, strains were first streaked on BHI medium (BD BBL). In each case, single colonies were subsequently plated both on CGXII-glucose (30) and CGXII-glutamine. The CGXII-glutamine broth is a modified version of medium CGXII that is devoid of (NH_4_)_2_SO_4_, urea and glucose and is supplemented with 100 mM glutamine. Plates were cultivated for 3 days at 30°C.

**Table 2.**
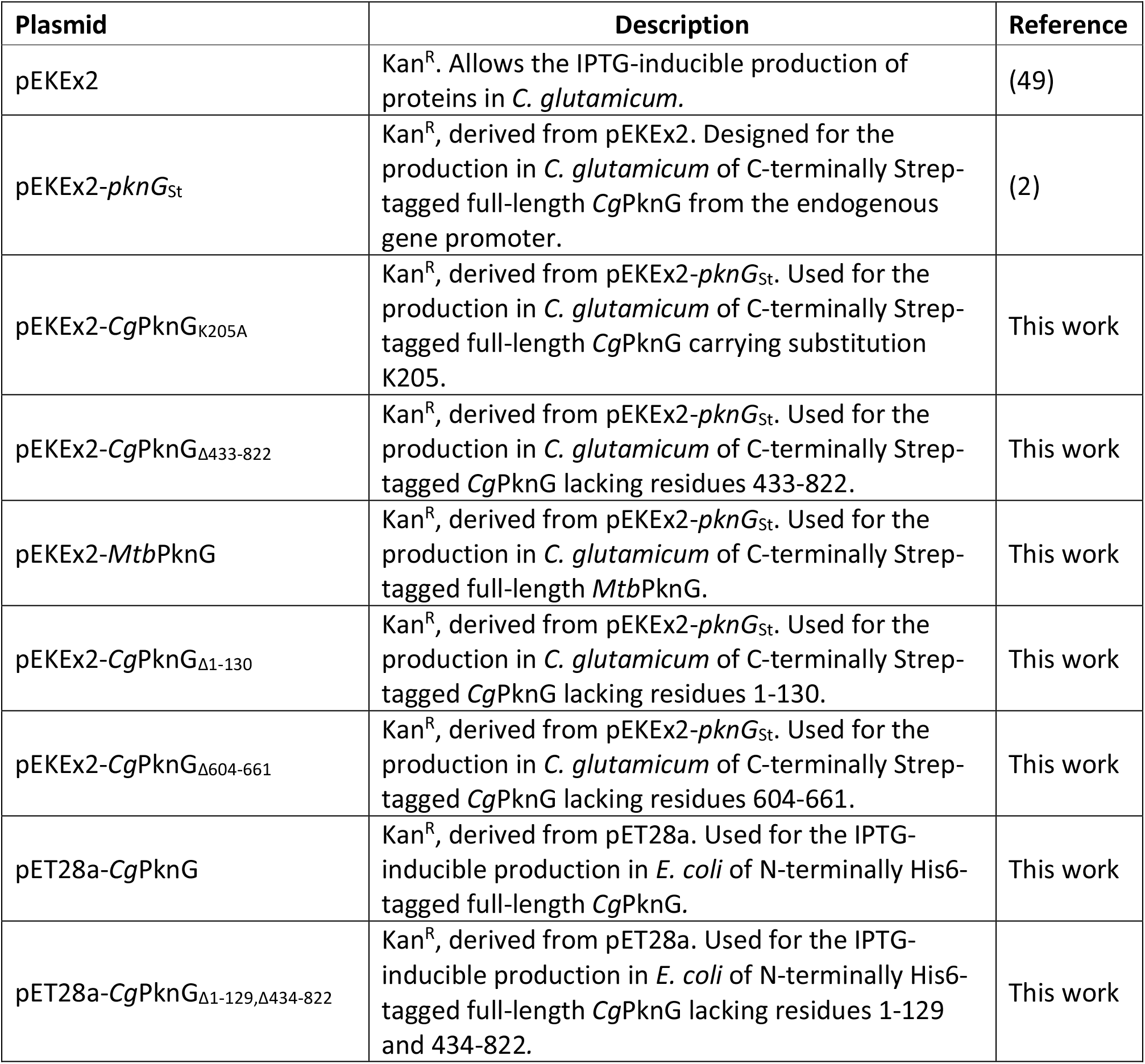
Plasmids used in this study.

### Detection of PknG versions by Western-blot

Transformed *C. glutamicum* Δ*pknG* (2) cells were grown at 30°C in BHI broth (BD BBL) with agitation until reaching 3 units of optical density at 600 nm. Protein expression was then induced by adding isopropyl β-D-1-thiogalactopyranoside (IPTG) to a final concentration of 1 mM, and the incubation was continued for 20 hours at 30°C. Cells were then harvested by centrifugation. Cell pellets were suspended in lysis buffer (50 mM Bis-Tris, 75 mM 6-aminocaproic acid, 1 mM MgSO_4_, 1 U/ml benzonase, cOmplete EDTA-free protease inhibitor cocktail (Roche) in the amount specified by the manufacturer, pH 7.4) and disrupted by using 0.1 mm glass beads and a homogenizer (Precellys 24) operated at 4°C. 120-250 μg of crude extracts were run in a pre-cast 4-12% SDS-PAGE gradient gel (Biorad) and then electro-transferred onto a 0.2 μm nitrocellulose membrane (Biorad). Blocking was performed with PBS buffer supplemented with 3% w/v BSA and 0.05% v/v Tween 20. The membrane was subsequently incubated with an anti-Strep antibody (StrepMAB-Classic, IBA Lifesciences) at 4°C overnight. After 3 washes with TBS-Tween buffer (10 mM Tris-HCl, 150 mM NaCl, 0.05 v/v Tween 20, pH 8.0) for 5 minutes each, the membrane was incubated with a secondary anti-Rabbit horseradish peroxidase conjugated antibody (GE Healthcare) for 45 minutes at room temperature. Finally, the membrane was washed 3 times with TBS-Tween buffer for 5 min each, revealed with the horseradish peroxidase substrate (Immobilon Forte, Millipore) and imaged using the ChemiDoc MP Imaging System (Biorad).

### Construction of plasmids for the production of recombinant proteins

Plasmids pET28a-*Cg*PknG and pET28a-*Cg*PknG_Δ1-129,Δ434-822_ (Table 2) were constructed by PCR amplification of *pknG* regions 1-822 and 130-433, respectively, from *C. glutamicum* ATCC 13032 genomic DNA, followed by digestion and ligation of the amplification products into the *Nde*I and *Sac*I sites in plasmid pET28a (Novagen). The oligonucleotides employed were the following (the TEV protease cleavage sites are underlined):

*Cg*PknG-F: ATTATCATATG<u>GAGAATCTTTATTTTCAGGGC</u>ATGAAGGATAATGAAGATTTCGATCC

*Cg*PknG-R: ATATTGAGCTCTCACTAGAACCAACTCAGTGGCCGCACGGC Δ1-129,Δ434-822-F:

TATATTATCATATG<u>GAGAATCTTTATTTTCAGGGC</u>GTTGCTGATGGCATGGTGGAATTG

Δ1-129,Δ434-822-R: TATATATTGAGCTCTCATTTGCCGTCGCGGACTGCCAAAATTTC

### Protein production and purification

Wild type *Cg*PknG and the truncation mutant *Cg*PknG_Δ1-129,Δ434-822_ were both overproduced in *E. coli* BL21(DE3) cells cultivated in LB broth. Wild type *Cg*PknG was produced for 18 h at 15°C with 500 µM IPTG, whereas *Cg*PknG_Δ1-129,Δ434-822_ was expressed after 3 h of induction at 30°C with 250 µM IPTG. Both of these proteins were then purified following the same protocol. *E. coli* cells were harvested by centrifugation, re-suspended in lysis buffer (25 mM Hepes, 500 mM NaCl, 20% v/v glycerol, 20 mM imidazole, pH 8.0), supplemented with cOmplete EDTA-free protease inhibitor cocktail (Roche) as specified by the manufacturer and sonicated. After clarification by centrifugation, the supernatant was loaded onto a HisTrap HP column (GE Healthcare) and the His-tagged protein was purified applying a linear imidazole gradient (20–500 mM) in lysis buffer. The His6-tag was later removed by over-night incubation at 4°C with 0.2 equivalents of His6-tagged TEV protease, followed by separation on a Ni-NTA agarose column (Qiagen). The protein was then further purified by size-exclusion chromatography on a 16/600 Superdex 200 column (GE Healthcare) equilibrated in either 50 mM Tris-HCl, 250 mM NaCl, 5% glycerol, pH 8.0 (wild type *Cg*PknG) or 25 mM Hepes, 150 mM NaCl, 5% glycerol, pH 7.5 (*Cg*PknG_Δ1-129,Δ434-822_), using a flow rate of 0.5-1 ml/min. Fractions corresponding to *Cg*PknG or *Cg*PknG_Δ1-129,Δ434-822_, as confirmed by SDS-PAGE, were pooled and concentrated, flash-frozen in liquid nitrogen and stored at -80°C.

GarA and OdhI were prepared as previously described (17, 31).

Proteins were quantified by using the molar absorption coefficient predicted from the aminoacid sequence by the ProtParam tool (http://web.expasy.org/protparam/).

### Protein kinase activity assays

Kinase activity assays were performed in 96-well plates. Each activity measurement was performed in a final volume of 20 µl, containing 50 mM Tris-HCl pH 7.4, 0.1% v/v 2-mercaptoethanol, 10 mM MnCl_2_, 100 µM [*γ*-^32^P]ATP (5-50 cpm/pmol), and 330 µM 17-mer peptide or 25 µM OdhI (or GarA) as substrate. The enzyme concentration in the assays was 0.7-3 µM and 0.15-0.9 µM when using the 17-mer peptide or OdhI (or GarA) as substrates, respectively. The kinase reactions were started by the addition of 4 µl [*γ*-^32^P]ATP-Mn^+2^ and were performed at room temperature. The reactions were stopped by the addition of phosphoric acid and 4 µl of each reaction were spotted on P81 phosphocellulose papers (Whatman) using the epMotion 5070 (Eppendorf) workstation. The papers were washed in 0.01% phosphoric acid, dried, then measured and analyzed using the PhosphorImager (FLA-9000 Starion, Fujifilm). Each reaction was performed in duplicates (<5% variation). In all cases, specific activity values were derived from reactions performed employing three different enzyme concentrations within the indicated ranges (<10% variation), verifying a linear dependence of activity with the enzyme concentration. Each assay was performed at least twice. The proportion of 17-mer peptide or OdhI (or GarA) phosphorylated in the reactions was lower than 10% and 30%, respectively. OdhI (or GarA) phosphorylation was verified to be linear in time up to 50% of its initial concentration. Under the experimental conditions employed to test phosphorylation of the 17-mer peptide or OdhI (or GarA), *Cg*PknG auto-phosphorylation represented less than 5% of the total signal. The measured signal was at least five times higher than the measure on the background.

The 17-mer peptide SDEVTV**ETTS**VFRADFL was produced with a purity >98% by Thermo Fisher Scientific.

### Mass spectrometry analysis

The kinase activity of *Cg*PknG was assayed using GarA as substrate and the molecular mass of unphosphorylated and phosphorylated GarA was then determined as previously described (15).

*Cg*PknG was incubated with ATP and MnCl_2_ and then sequentially digested with trypsin and endoproteinase GluC for 3 h at 37°C. The resulting peptides were separated using a nano-HPLC system (Proxeon EasynLC, Thermo) with a reverse-phase column (easy C18 column, 3 μm; 75 μm ID×10 cm; Proxeon, Thermo) and eluted with a 0.1% v/v formic acid (in water) to acetonitrile gradient (0–40% acetonitrile in 50 min; flow 300 nl/min). Online MS analysis was carried out in a linear ion trap instrument (LTQ Velos,Thermo) in data dependent acquisition mode (full scan followed by MS/MS of the top 5 peaks in each segment, using a dynamic exclusion list). Raw MS/MS spectra were extracted by the Proteome Discoverer software package (v.1.3.0.339, Thermo) and submitted to Sequest for database searching against sequences from *E. coli* (strain K12) downloaded from Uniprot consortium (April, 2021) to which the sequence of PknG from *C. glutamicum* was added. Search parameters were set as follows: peptide tolerance: 0.8 Da; MS/MS tolerance: 0.8 Da; methionine oxidation and Ser/Thr/Tyr phosphorylation as the allowed variable modifications. PhosphoRS was used as phospho-site localization tool (32). We considered a positive phospho-site identification when more than one spectrum for the phospho-peptide was obtained, pRS probability was >95% and manual inspection of the MS/MS spectra showed at least two confirmatory fragment ions.

### Crystallization and data collection

Crystallization screenings were carried out using the sitting-drop vapor diffusion method and a Mosquito nanolitre-dispensing crystallization robot (TTP Labtech). Crystals of *Cg*PknG_ΔN-t_ + AMP-PNP and *Cg*PknG_Δ1-129,Δ434-822_ + AMP-PNP grew after 20-30 and 7-10 days, respectively, from 10 mg/ml protein solutions supplemented with 5 mM AMP-PNP, by mixing 200 nl of protein solution and 200 nl of mother liquor (100 mM Tris-HCl, 17% w/v PEG 20 k, 100 mM MgCl_2_, pH 8.5; and 100 mM Tris-HCl, 27-30% w/v PEG 4 k, 200 mM MgCl_2_, pH 8.8, respectively), at 18°C. Single crystals reaching a size of (100 µm)^3^ were cryprotected in mother liquor containing 25% glycerol and flash-frozen in liquid nitrogen. X-ray diffraction data were collected at the synchrotron beamlines Proxima 2 (Synchrotron Soleil, Saint-Aubin, France) and ID29 (ESRF, Grenoble, France) at 100 K. Employed wavelengths were 0.9801 Å and 0.97625 Å for *Cg*PknG_ΔN-t_ + AMP-PNP and *Cg*PknG_Δ1-129,Δ434-822_ + AMP-PNP crystals, respectively. The diffraction data were processed using XDS (33) and scaled with Aimless (34) from the CCP4 program suite.

### Structure determination and refinement

The crystal structure of *Cg*PknG_ΔN-t_ + AMP-PNP was solved by molecular replacement using the program Phaser (35) and the atomic coordinates of *Mtb*PknG residues 138-405 from PDB 4Y0X (13) and residues 406-750 from PDB 2PZI (12) as search probes. The structures of *Cg*PknG_Δ1-129,Δ434-822_ + AMP-PNP were solved similarly by using the atomic coordinates of *Cg*PknG_ΔN-t_ residues 165-425. Ligand molecules were manually placed in *mFo–DFc* sigma-A-weighted electron density maps employing *Coot* (36). Models were refined through iterative cycles of manual model building with *Coot* and reciprocal space refinement with phenix.refine (37). The final models were validated through the MolProbity server (38). In each case, the final model contained more than 97% of residues within favored regions of the Ramachandran plot, with no outliers. Figures were generated and rendered with Pymol 1.8.x. (Schrödinger, LLC).

### Edman degradation

The crystal employed to solve the structure of *Cg*PknG_ΔN-t_ was dissolved in water and Edman degradation was performed by the Functional Genomics Center of Zurich (https://fgcz.ch/omics_areas/prot/applications/protein-characterization.html). As a control, an aliquot of recombinant *Cg*PknG as used in crystallization screenings was also analyzed, and the sequence of the protein N-terminus resulted GMKDN, as expected.

### Analytical ultracentrifugation

Sedimentation velocity experiments were carried out at 20°C in an XL-I analytical ultracentrifuge (Beckman Coulter). Samples were spun using an An60Ti rotor and 12-mm double sector epoxy centerpieces. The partial specific volume of *Cg*PknG (0.734 ml g^−1^) was estimated from their amino acid sequences using the software Sednterp. The same software was used to estimate the buffer viscosity (η = 1.040 centipoises) and density (ρ = 1.010 g·ml^−1^). *Cg*PknG (400 μl at 1 mg/ml) was spun at 42,000 rpm, and absorbance profiles were recorded every five minutes. Sedimentation coefficient distributions, c(s), were determined using the software Sedfit 14.1 (39).

### Database searches, alignments and phylogenetic analyses

BLASTp searches (40) were conducted against complete protein sequences available at the Integrated Microbial Genome (IMG; http://img.jgi.doe.gov) (41), performing a taxon sampling on finished assembled genomes within the phyla *Cyanobacteria, Chloroflexi, Chlorobi, Fusobacteria, Sinergistetes, Firmicutes, Tenericutes, Acidobacteria, Nitrospirae, Spirochaetes, Aquificae* and *Thermotogae*, all in the vicinity of *Actinobacteria* in an updated tree of life (27). The sequence of *Mtb*PknG was used as queries for searches to identify homologues in such genomes using an expected inclusion threshold e-value < 1 e^-20^. Once the existence of the domain combinations was confirmed, we focused on 91 complete *Actinobacteria* genomes available from IMG (April 2021). The final selection was preprocessed using PREQUAL (42) to mask non-homologous sequence stretches. A CD-HIT (43) cut-off value of 90% pairwise identity was applied for the entire set of sequences retrieved as described. The final set of 40 sequences was aligned with MAFFT (version 7.467) using the L-INS-I strategy (44) and columns with more than 90% gaps were removed with TrimAL. The phylogenetic tree displayed in Fig. S7 was computed with IQ-TREE (version 1.6.12, (45)) using ModelFinder (46) to select the evolutionary model and the ultrafast bootstrap method (47) (options “-bb 1000 -alrt 1000”). The model selected with the Bayesian Information Criterion was the evolutionary matrix EX_EHO (48) with empirical frequencies and four categories of free rate (EX_EHO+F+R4).

### Data availability

Atomic coordinates and structure factors have been deposited in the Protein Data Bank under the accession codes 7mxb (*Cg*PknG_ΔN-t_ + AMP-PNP), 7mxj (*Cg*PknG_Δ1-129,Δ434-822_ + AMP-PNP_1) and 7mxk (*Cg*PknG_Δ1-129,Δ434-822_ + AMP-PNP_2).

## ACKNOWLEDGEMENTS

MNL received fellowships from the EMBO (European Molecular Biology Organization) and the Fondation pour la Recherche Médicale (FRM, France). This work was funded by grants from the Institut Pasteur, the CNRS (France), the Agence Nationale de la Recherche (ANR, France, contract number ANR-09-BLAN-0400) and the European Commission Seventh Framework Programme (contract HEALTH-F3-2011-260872). MNL acknowledges support from the Agencia I+D+I (grant PICT 2017-1932). RMB acknowledges support from DFG (grant BI 1044/12-1).

We thank Ahmed Haouz and Patrick Weber for their help with robot-driven crystallization screenings and Bertrand Raynal for his help with analytical ultracentrifugation experiments. We acknowledge William Shepard for his assistance in the collection of diffraction data at beamline Proxima 2 and Martin Cohen-Gonsaud for providing plasmid pET-15b-Tev-OdhI for OdhI production.

## Author contributions

MNL designed experiments, prepared proteins, performed kinase activity assays, carried out crystallographic studies and structural analysis, analyzed data and wrote the paper; ASC performed complementation assays, analyzed data and wrote the paper; NB prepared plasmids pET28a-*Cg*PknG and pET28a-*Cg*PknG_Δ1-129,Δ434-822_, optimized the production of recombinant proteins and performed analytical ultracentrifugation experiments; MGi carried out mass spectrometry analyses; MGa performed phylogenetic analysis; RD designed and performed mass spectrometry studies; RMB designed kinase activity assays and analyzed data; MBe, MBo and PMA designed research and analyzed data. All authors copy edited the paper.

